# SARS-CoV-2 (E)-protein induces rapid TLR2-mediated T cell activation in mouse lungs revealed by intravital lung microscopy

**DOI:** 10.64898/2026.05.03.722459

**Authors:** Yasmin Shaalan, Nethaji Kuruppu, Zane Orinska, Chenxi Li, Frauke Koops, Roxana-Maria Wasnick, Elfriede Noessner, Tobias Stoeger, Silke Meiners, Markus Rehberg

## Abstract

Mounting evidence indicates that T cells can operate in an innate-like mode challenging the classical description of T cells as strictly adaptive immune effectors. T cells can engage innate pattern recognition receptors to mount rapid but antigen-nonspecific responses to infection or cellular stress. This study observed that CD8+ T cells, and to a lesser extent also CD4+ T cells, responded to viral proteins in the mouse lung quickly in an innate-like fashion.

We employed intravital lung microscopy to visualize infiltration of CD8+ T cells into the lung following intratracheal instillation of the SARS-CoV-2 envelope (E)-protein. Here, we demonstrate acute recruitment of CD8+ from the pulmonary microcirculation into the lung as early as 4 and 24 hours after (E)-protein instillation. The acute infiltration of CD8+ T cells was not observed in Tlr2^−^/^−^ mice. Immunohistochemistry analysis of mouse lungs revealed T cell accumulation in nodular inflammatory foci (NIF) of the lung at perivascular regions and around large airways. Stimulating spleen-derived CD8+ T cells from wild-type mice with (E)-protein *ex vivo* in combination with cytokines or TCR agonists significantly upregulated CD69 and activated secretion of interferon (IFN)γ which was not observed with CD8+ T cells isolated from Tlr2^−^/^−^ mice.

These findings indicate rapid bystander activation of CD8+ T cells by the SARS-CoV-2 envelope (E)-protein that depends on (E)-protein sensing by TLR2. This innate-like CD8+ T cell response to SARS-CoV-2 (E)-protein may offer novel opportunities for diagnostic and therapeutic development, warranting further investigation.

## Introduction

Respiratory viruses such as severe acute respiratory syndrome coronavirus 2 (SARS-CoV-2) have caused major public health concerns, with hundreds of millions of reported cases worldwide (World Health Organization, 2026). Since the lung is continuously exposed to environmental antigens and pathogens, pulmonary immunity needs to balance the generation of rapid and effective antiviral responses while maintaining tissue integrity and homeostasis. Failure to achieve this balance can result in excessive inflammation, tissue damage, and impaired gas exchange (Kohlmeier & Woodland, 2009).

The SARS-CoV-2 envelope (E)-protein is a structural protein of the coronavirus family and an important determinant of its viral pathogenicity (Schoeman & Fielding, 2019). Its conservation across SARS-CoV-2 variants and related coronaviruses makes it an attractive candidate for therapeutic and preventive strategies (Alam et al., 2020). In addition to its structural role, the (E)-protein acts as a viroporin that disrupts host membrane integrity and contributes to cell injury and death (Planès et al., 2021).

In the respiratory tract, viral infections are detected by pattern recognition receptors (PRRs) expressed by immune cells such as alveolar macrophages, dendritic cells, monocytes, neutrophils, NK cells, B cells, and T cells, as well as non-immune cells including epithelial and endothelial cells and fibroblasts (Vijay, 2018; Zarember & Godowski, 2002). PRR engagement activates signalling pathways involving NF-κB and interferon regulatory factors (IRFs), which induce the release of chemokines and proinflammatory cytokines such as IL-6, TNFα, CCL2, and type I interferons (J. P. Wang et al., 2007). These mediators are essential for antiviral defence but can also contribute to immunopathology if produced excessively or for prolonged periods.

Among PRRs, TLR2 has emerged as an important sensor of β-coronaviruses. SARS-CoV-2 surface proteins can activate TLR2 independently of viral replication, triggering ERK and NF- κB signalling in macrophages and inducing expression of inflammatory genes such as IL1B, IL6, and TNF (Mantovani et al., 2023; Zheng et al., 2021). In the mouse and human system, TLR2 inhibition or deletion reduces cytokine production and dampens inflammatory responses to SARS-CoV-2, supporting a central role for this pathway in coronavirus-induced inflammation (Zheng et al., 2021). These findings suggest that innate sensing of the (E)-protein can occur before productive infection and may help initiate early immune activation in the lung.

Adaptive immune responses are classically initiated when naïve T cells receive three coordinated signals: T cell receptor (TCR) recognition of peptide-MHC complexes, co-stimulation through CD28-CD80/CD86 interactions, and cytokine-mediated polarization (Curtsinger & Mescher, 2010). However, accumulating evidence shows that T cells can also be activated in a TCR-independent manner during infection or inflammation, a process known as bystander activation. In this context, inflammatory cytokines or innate receptor signals can induce effector-like functions in T cells without cognate antigen recognition, challenging the classical view that T-cell activation is exclusively antigen-specific (Yosri et al., 2024). Bystander activation is particularly relevant for CD8+ T cells, which can rapidly acquire effector functions such as IFNγ production, granzyme expression, and NKG2D upregulation in response to cytokines including IL-12, IL-15, IL-18, and type I interferons. This response can enhance early antiviral defence, but it may also contribute to tissue injury and sustained inflammation if excessive or poorly controlled (Crosby et al., 2014; Maurice et al., 2021).

Here, we show that SARS-CoV-2 E protein induces rapid CD8+ T cell activation *ex vivo* and *in vivo* in a TLR2-dependent manner. This mechanism may represent an important bridge between innate recognition of a viral structural protein and rapid functional reprogramming of CD8+ T cells.

## Material and Methods

### Animals

Wildtype C57BL/6J mice were purchased from Charles River Laboratories (Sulzfeld, Germany). Tlr2^-/-^ (B6.129-*Tlr2*^*tm1Kir*^*/J*) mice were kindly provided by Carsten Kirschning, Institute of Medical Microbiology, University Hospital Essen, Germany. Mice were housed in accordance with Helmholtz Zentrum Muenchen institutional, state and federal guidelines. Experiments were performed according to the guidelines of the Regierung von Oberbayern (District Government of Upper Bavaria) under the approval number ROB-55.2Vet-2532.Vet_02-19-150. Also, mice were bred under specific pathogen-free (SPF) conditions at the Research Center Borstel animal facility. During experiments, animals were housed in type II long IVCs with wood bedding (Abedd Espe Classic 2-5mm LTE E-001) at 20-24°C, 45-65% relative humidity, and a 12h light/dark cycle. γ-Irradiated maintenance diet (ssniff Spezialdiäten GmbH, Soest) and water were provided *ad libitum*. Each cage contained nesting material and red plastic boxes for environmental enrichment. Mice of both sexes were used for experiments.

### Lung-intravital microscopy

Recombinant SARS Cov2 (E)-protein was instilled at a dose of 25 µg per mouse (100-200 days old). Four- or 24-hours post-instillation, lung intra-vital microscopy (lung-IVM) was performed as described in detail in (Liu et al., 2025). Mice were first anesthetized by the intraperitoneal injection of medetomidine (0.5 μg/g), midazolam (5 μg/g) and fentanyl (0.05 μg/g). Fluorescent antibodies (anti-CD4-PE, 2 μg/mouse (clone GK1.5), anti-CD8-APC, 3 μg/mouse (clone 53-6.7), both from BD Pharmingen™) and anti-Ly6G-AF488, 3 μg/mouse (clone 1A8, BioLegend)), were injected 20 min before performing thoracic surgery, vacuum stabilization of the lungs and fluorescence microscopy. Time lapse images were acquired by VisiScope A1 imaging system (Visitron Systems GmbH) for 10 min in 5 different alveolar regions per mouse. The vital signs of the mouse were regularly checked throughout the acquisition time.

### Immunofluorescence of lung tissue

Mice were treated with (E)-protein for 24 hours, followed by euthanization using ketamine (0.5 μg/g). Lungs were intratracheally filled with 4% paraformaldehyde (PFA) in PBS, stitched and stored inverted overnight in 4% PFA. The next day, PFA was discarded and each lobe was cut into thin slices and transferred to tissue embedding cassettes and processed to generate lung sections of 3 μm thickness by Epredia™Section Transfer System™ for rotary microtome from Fischer Scientific, which were then transferred to microscopic slides. Staining of the sections were done after standard deparaffinization, antigen retrieval, blocking with goat serum (1 hour, room temperature (RT)). All washing steps were performed in PBS containing 0,05% Tween2. Following rabbit antibodies were used as primary antibodies for sequential staining at 4°C overnight: anti-CD8α (clone EPR20305, Abcam), anti-MHC Class II (I-Ab) (PA5-116876), anti-NKG2D (PA5-97904) - both from Thermo Fischer Scientific and anti-GzmB (clone D6E9W, Cell Signaling Technology). After incubation and washing slides were incubated with anti-rabbit secondary antibodies at dilution of 1:1000 for 1 hour at room temperature. Slides were incubated in rabbit blocking serum for 1 hour at room temperature, washed and blocked with Fab fragment (Goat Fab anti-Rabbit IgG) diluted in goat blocking serum at 1:45 for 90 min in 37°C. Slides were washed and incubated with the next primary antibody at 4°C overnight, washed and incubated with the second secondary antibody as well as DAPI at a dilution of 1:1000 for 1 hr at room temperature. Slides were then washed, mounted and imaged with Zeiss Axiovert Inverted Fluorescence Microscope (Carl Zeiss). For each mouse lung, 10 to 20 images from randomly selected areas were saved for further analysis. Fiji image J software was used for image analysis.

### BAL Analysis

Bronchoalveolar lavage (BAL) fluid and cells were collected by cannulating the trachea and rinsing the lung four times with PBS. Cells were collected for cytospin analysis. The lavage fluid and mouse serum were used to perform a cytokine array.

For the cytospins, 3×10^4^ cells were put on a slide with a cyto-funnel, centrifuged at 20xg for 6 min using Cytospin 2 Cytocentrifuge (Shandon). Cytospin slides were then dried and stained in May Grunwald and Giemsa stain and examined as described before (Voss et al., 2025).

### Cytokine analysis

Bio-Plex Pro Mouse Chemokine Panel, 31-plex Assay from Bio-Rad was used to detect and quantify chemokines. BAL fluid and serum samples were applied to the plates closely following the manufacturer’s instructions. The plate was read on a Bio-plex plate reader. Instrument settings were adjusted according to the manual instructions.

### Flow cytometry analysis of lung cells

Lungs were dissociated into single cell suspension using Miltenyi Biotec mouse lung dissociation kit, in conjunction with gentle MACS Octo dissociator (Miltenyi Biotec). Digestion was followed by filtration through a 70 μm SmartStrainer filter, followed by centrifugation at 300xg for 10 min. Ery-lysis buffer (150 mM NH_4_Cl, 10 mM KHCO_3_, 500 nM EDTA, pH 7.2) was applied for a 2 min, followed by washing with flow cytometry buffer (0.5% FBS, 2 mM EDTA, PBS).

Each sample was resuspended in 100 μl of flow cytometry buffer. Fluorochrome-conjugated antibodies from Miltenyi Biotec (anti-CD45 PerCp Vio700 (clone REA737), anti-CD3 APC (clone REA606), anti-CD4 PE (clone REA604), anti-CD8 FITC (clone REA734), anti-MHC class II PE-Vio 615 (clone REA813)) were used for staining at a dilution of 1:50. Unstained, single stained and Fluorescence Minus One (FMO) controls were included for all antibodies to ensure choosing the right voltage, compensation and gating. All samples were shortly vortexed, and staining was done for 15-20 min at 4°C, protected from light. Sample was then washed twice in flow cytometry buffer and centrifuged for 5 min 400xg at 4°C. Samples were resuspended in 400 μl of flow cytometry buffer. Cells from lung tissue were re-filtered into Falcon™ Round-Bottom Polystyrene Test Tubes with Cell Strainer Snap before measurements. DAPI was added for dead cell exclusion. Flow cytometry data were collected using BD FACSCelesta™ flow cytometer and analyzed using FlowJo (v.10) software.

### CD8+ T cell stimulation in vitro

96-well plates (Corning) were coated with antibodies anti-CD3 (1 μg/ml, clone 145-2C11, eBioscience) and anti-CD28 (5μg/ml, clone 37.51, BD Pharmingen™) in PBS overnight at 4°C. The following day, C57Bl/6*J* and Tlr2^-/-^ mice were euthanized and blood samples (in 2.7% EDTA in PBS) and spleens were collected. Blood leukocytes were isolated by lysis of red blood cells in lysis buffer I (150 mM NH_4_Cl, 10 mM Na_2_CO_3_, 1 mM EDTA, pH 7.2). CD8+ T cells from spleens were isolated by disassociating the tissue using a syringe plunger, erythrocytes were lysed with lysis buffer II (150 mM NH_4_Cl, 10 mM KHCO_3_, 0.1 mM EDTA, pH 7.2) for 2 min on ice and cell suspension passed through a 30 μm CellTrics™ cell strainer. The cell suspension was centrifuged at 427xg for 5 min at 4°C. Isolation of CD8+ T-cells was done using the mouse CD8+ isolation kit (Miltenyi Biotec) according to the manufacturer’s instructions. Cell purity was determined by staining with antibodies in staining buffer (2% FBS, 4 µM EDTA, 0.1% NaN_3_ in PBS) and propidium iodide (PI, 1 µg/ml in staining buffer, Sigma) and measured in the MACSQuant Analyzer 16 (Miltenyi Biotec). Cells were washed in cell culture media (RPMI 1640, 10% FCS, 10 U/ml penicillin, 0.1 mg/ml streptomycin, 2 mM L-glutamine, 50 µM 2β- mercaptoethanol) and plated 1×10^6^ cells/ml either without any stimulation (control) or with addition of (E)-protein (2 μg/ml, Thermo Fisher Scientific), Pam3CSK4 (10 ng/ml, EMC microcollections), together with cytokines (interleukin-2 (IL-2, 20 ng/ml, Biotest) and IL-15 (5 ng/ml R&D Systems)) or plate-coated antibodies and incubated at 37°C, 5% CO_2_ for 16 hr. Plates were centrifuged at 706xg for 3 min at 4°C, supernatants and cells were collected.

### Flow cytometry of cultured cells

CD8+ T cell activation was determined by staining with antibodies (anti-CD69 BV421 (clone H1.2F3), anti-TCRβ FITC (clone H57-597), anti-I-A/E PE (clone M5/114.15.2), anti-CD8α APC (clone 53-6.7) (all from Biolegend), anti-CD314 PE-Cy7 (clone CX5, eBioscience), in staining buffer and detected by MACSQuant Analyzer 16. Dead cells were excluded by PI staining. Flow cytometry files were analysed with FlowLogic software (v.8, Inivai Technologies) and statistical analysis was performed in GaphPad Prism (v.10).

### Analysis of IFNγ production

IFNγ production in supernatants of stimulated WT and Tlr2^-/-^ CD8+ T were analyzed using the U-PLEX Mouse IFNγ Assay from MSD according to manufacturer’s instructions and measured in MESO QuickPlex SQ 120MM.

## Results

### (E)-protein induces rapid CD8+ T cell recruitment into the lung

To investigate the effects of SARS-CoV-2 (E)-protein on lung T cells, intravital lung microscopy (lung-IVM) was employed. Lung-IVM was performed 4 or 24 hours after intratracheal instillation of recombinant (E)-protein and circulating lymphocytes were visualized by labelling CD8+ and CD4+ T cells (Fig 1A). This *in vivo* labelling approach uncovered acute infiltration of CD8+ T cells from the vasculature into the perivascular region of the lung at 4 hours post instillation (Fig. 1B), as also confirmed by flow cytometry in whole lungs at 4 hours (Fig. 1C). After 24 hours, this response was still detectable in lung-IVM but no longer in the flow cytometry analysis (Fig. 1 C). This difference might be because flow cytometry detects all CD8+ T cells in the lung including infiltrating and resident T cells while in the *in vivo* imaging experiments intravenous injection of fluorescently labelled antibodies was used to specifically detect CD8+ T cells infiltrating the lung from the vasculature. Notably, no significant recruitment of CD4+ T cells was observed by intravital microscopy (Fig. 1B).

**Fig. 1.**
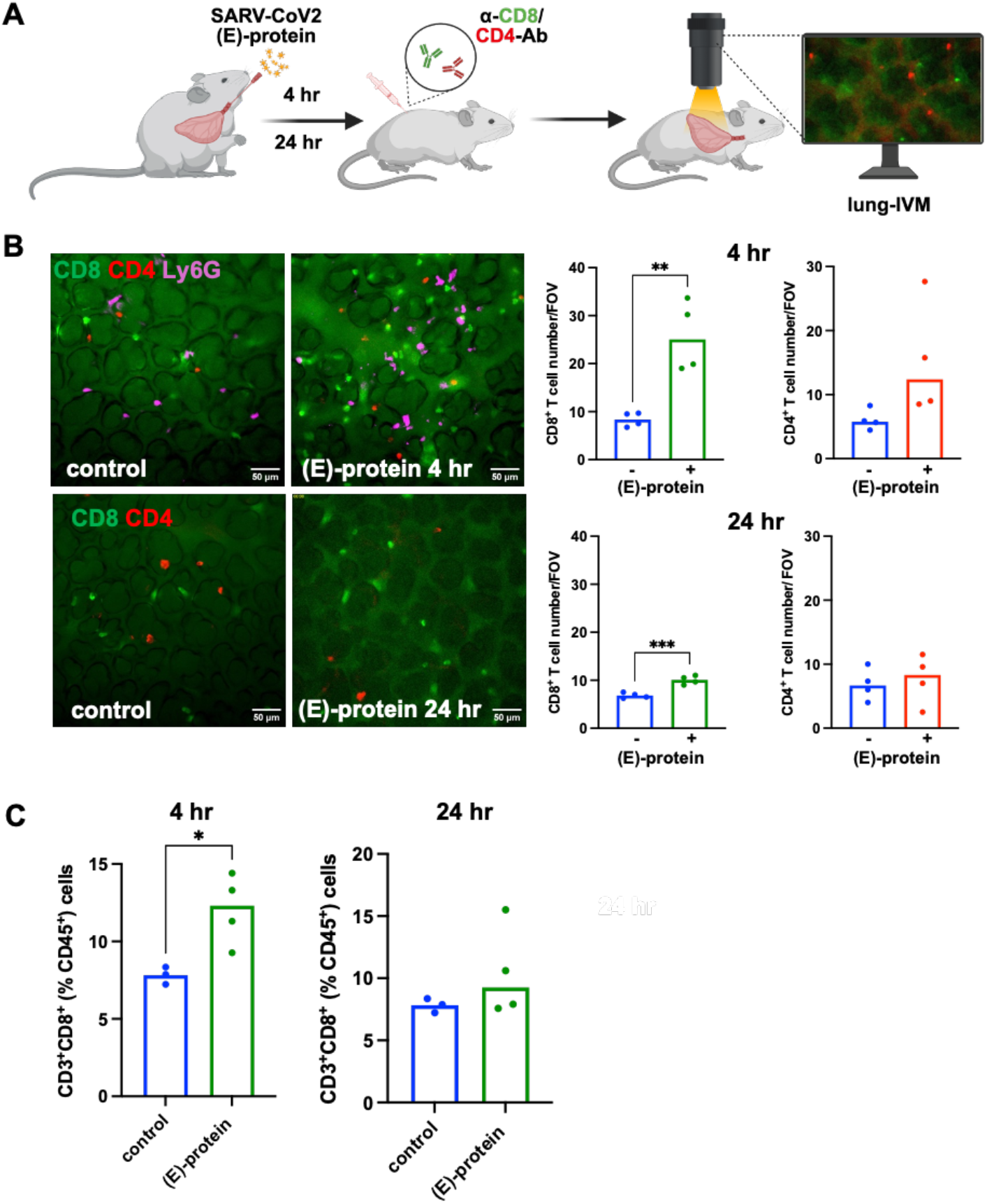
Intravital lung microscopy (lung-IVM) reveals fast recruitment and activation of CD8+ T cells upon challenge with SARS-CoV2 (E)-protein. **A)** Experimental scheme: C57Bl/6J mice were instilled with 25 μg/mouse (E)-protein and investigated after 4 or 24 hours (hr). T cells were visualized via intravenous injection of fluorescently labelled antibodies against CD8 or CD4 20 minutes (min) before performing lung-IVM. **B)** Representative images taken from lung-IVM time lapse videos (duration 10 min) are shown with CD8+ T cells labeled in green and CD4+ T cells labeled in red after 4 and 24 hr after (E)-protein instillation. Granulocytes were labelled using Ly6G-AF488 (3 μg/mouse). CD8+ and CD4+ T cells were quantified per Field of View (FOV, 0.16 mm^2^) using the mean of 5 FOV/mouse (n=4 mice/group). Contrast rich roundish structures in the lung tissue, outlined by tissue autofluorescence in the green channel, are alveoli, surrounded by microvessels. Unpaired t-test was applied using the respective experimental control with p<0.001 (***) and p<0.01 (**). **C)** Percentage of CD3+CD8+ T cells of CD45+ cells in lungs of mice that had been instilled with (E)-protein 4 hr and 24 hr before. Unpaired t-test was applied with p<0.05 (*).

### (E)-protein triggers neutrophil influx and pro-inflammatory cytokine release

Cytospin analysis of BAL fluid uncovered an increased recruitment of neutrophils into the alveolar space, particularly after 24 hours of (E)-protein instillation while we did not detect significant recruitment of other immune cells like macrophages and lymphocytes within 24 hours (Fig. 2A). We next analysed a panel of cytokines in BALF and blood serum to identify potential signalling pathways that would contribute to the acute infiltration of T cells into the lung after the (E)-protein challenge. We observed a distinct cytokine profile in BALF and blood that was specifically regulated after 4 and 24 hours upon (E)-protein instillation (Fig. 2B). Induction of inflammatory cytokines peaked at 4 hours in the blood suggesting a rapid systemic sensing of (E)-protein. The cytokine changes in the BALF were less prominent after 4 hours but more pronounced after 24 hours except for some specific cytokines such as TNFα, IL6, CXCL1, CCL3, 4, 20 (Fig. 2B).

**Figure 2.**
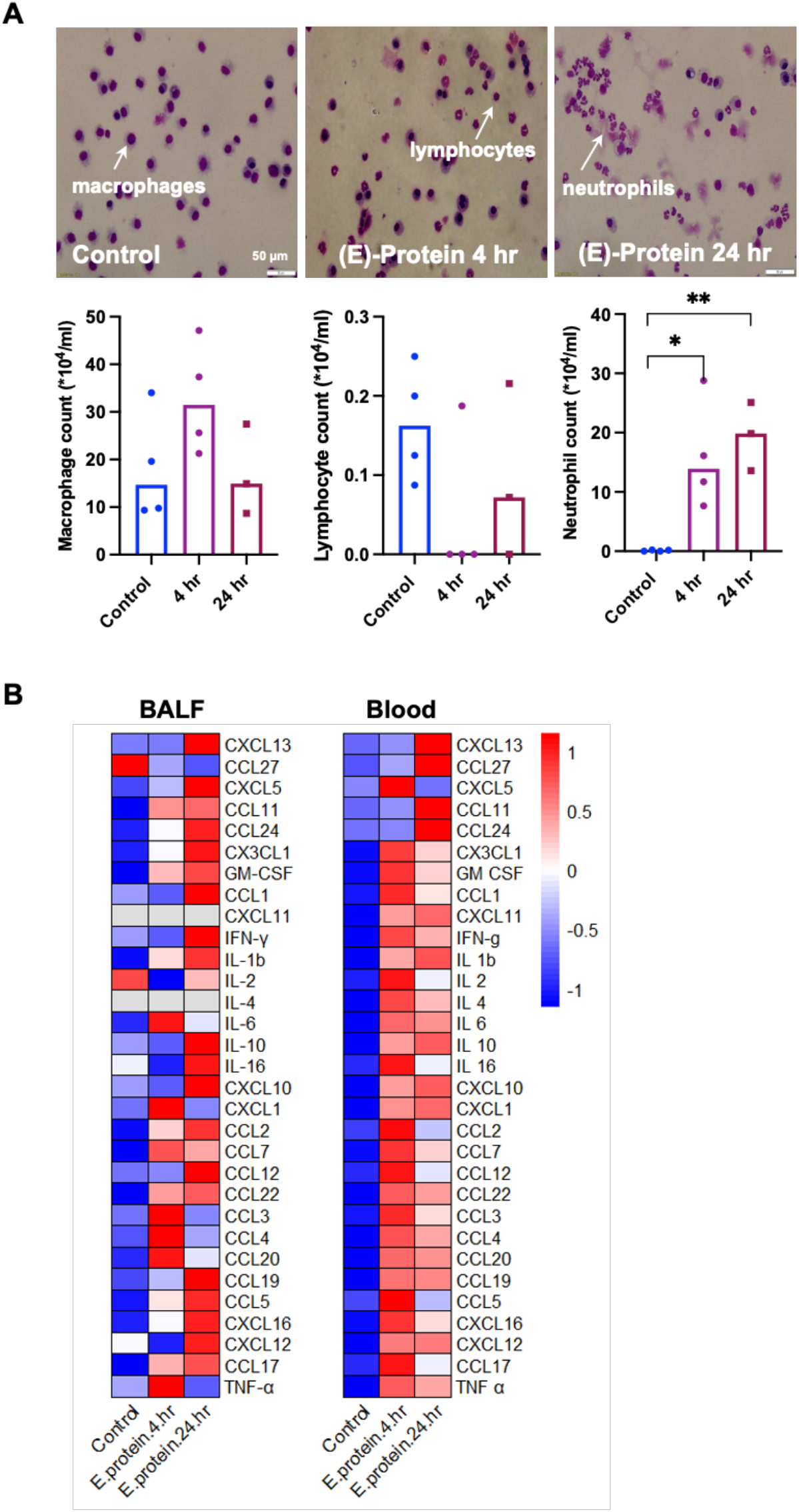
BAL analysis shows neutrophilia and elevated proinflammatory response upon (E)-protein stimulation. **A)** May-Grünwald-Giemsa staining of bronchoalveolar lavage fluid (BAL). Mice were treated with (E)-protein and analyzed after 4 or 24 hr. Identification of cells was based on differences in cellular morphology and colour. Corresponding quantification of neutrophil, lymphocytes and macrophages counts in BAL. One-way Anova was applied with p<0.01 (**) and p<0.05 (*). **B)** Heatmap showing the concentration of 29 cytokines in bronchoalveolar lavage fluid (BALF) and blood serum of mice in pg/ml. Mice were instilled with (E)-protein and samples were collected 4 and 24 hr post instillation. Each data point is the mean of cytokine concentrations obtained from 4 different mice per treatment group.

### T cells accumulate in nodular inflammatory foci

To characterize CD8+ T cell recruitment patterns in microvessels of the alveolar region, detailed time-lapse analysis of the CD8+ T cell dynamics was performed and revealed a prominent decline in migration velocity following (E)-protein instillation (Fig. 3A). While in the control group T cells exhibited transient contacts with microvascular walls (“vessel-tethering behavior”), (E)-protein treatment induced increasing numbers of T cells to maintain prolonged contacts. Specifically, the majority of these CD8 + T cells transitioned to a crawling phenotype within the alveolar region, a behaviour that persisted for up to 24 hours (Fig. 3B). Immunofluorescent analysis of lung tissue co-stained for CD8+ T cells and MHC class II^+^ antigen presenting cells (APC) revealed the accumulation and aggregation of T cells and APCs in nodular inflammatory foci (NIF) enriched at perivascular regions and around large airways of the lung in (E)-protein challenged mice compared to control tissue (Fig. 3C). Such NIFs, characterized as organized clusters of APCs and T cells that form at sites of infection or inflammation, e.g. in the lung, are thought to orchestrate local immune responses (Stahl et al., 2013). In addition, we observed that CD8+ T cells within these NIFs prominently expressed the innate activation marker NKG2D and the cytotoxic effector molecule granzyme B (Fig. 3D). Taken together, our lung-IVM, flow cytometry and immunofluorescence stainings suggest that pulmonary delivery of SARS-CoV2 (E)-protein triggers rapid recruitment of CD8+ T cells into the lungs followed by their assembly into localized inflammatory structures and differentiation into activated cytotoxic effector T cells.

**Fig. 3.**
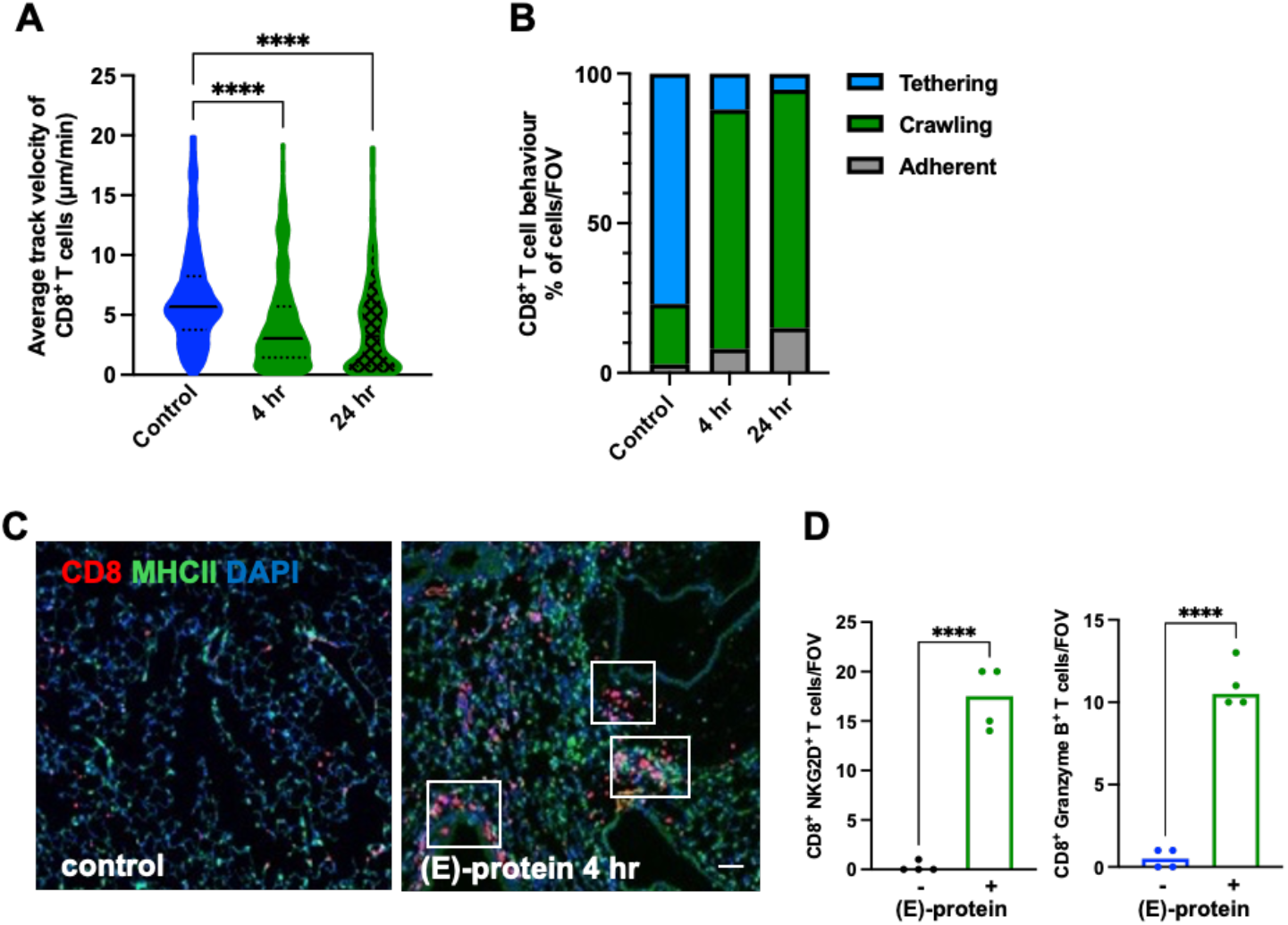
T cells accumulate in nodular inflammatory foci (NIF). **A)** Average track velocity and **B)** classification of CD8+ T cell behaviour into crawling, tethering or adherent CD8+ T cells in control mice and mice instilled with (E)-protein 4 or 24 hr before. Each data point represents an individual cell. Adherent cells were defined as cells that showed no displacement during the 20 min of imaging, crawling cells are T cells with an average velocity of less than 20 µm/min, while tethering cells are defined as cells that jump fast from one point to the next within the Field of View (FOV, 0.16 mm^2^) using Fiji Image J software (TrackMate plugin). One-way ANOVA with Dunnett’s multiple comparison testing was applied with p<0.0001 (****). For velocity measurements, 500-1000 cells were analyzed, behaviour classification was done with 100 CD8+ T cells in total per condition. **C)** Representative whole lung sections showing nodular inflammatory foci (NIFs) in lungs of mice that had been intratracheally instilled with (E)-protein 24 hr before. FFPE section of lungs were stained for CD8 and MHC class II to identify CD8+ T and antigen presenting cells, respectively, with DAPI (Sigma Aldrich) as a nuclear stain to detect all cells. Bar represents 50 µm. **D)** Quantification of immunofluorescence staining of CD8+ T cells expressing NKG2D and GzmB in paraffin embedded lungs of mice that had been intratracheally instilled with (E)-protein 24 hr before. CD8+ NKG2D+ and CD8+ GzmB+ T cells were quantified per FOV (FOV, 0.23 mm^2^). 5 FOV/mice were counted, and the mean was calculated for each mouse (n=4/group). Unpaired t-test was applied with p<0.0001 (****).

### (E)-protein activates CD8 + T cells via TLR2-dependent bystander mechanisms ex vivo

As these mice were not previously exposed to SARS-CoV2 (E)-protein, we hypothesized that this rapid CD8+ T cell activation might occur through a mechanism akin to bystander activation, which is defined as antigen-independent activation driven by inflammatory cues, particularly cytokines. Co-stimulation classically refers to a second signal required following antigen-specific TCR engagement (Lee et al., 2022).

To explore this, we isolated CD8+ T cells from splenocytes of naïve mice and stimulated them with (E)-protein, and in parallel with cytokines and TCR agonists (αCD3/αCD28 antibodies), and the combination of both. Both cytokine and antibody treatments yielded slightly better cell viabilities compared to the unstimulated control, demonstrating that co-stimulation is favoured by T cells (Fig. 4A, B). (E)-protein together with cytokines or TCR agonists significantly upregulated the expression of CD69 on isolated CD8+ T cells but was ineffective by itself (Fig. 4C, D). Similar effects were observed with the TLR2 agonist Pam3CSK4. The effect of (E)- protein on CD69 co-activation was absent in CD8+ T cells isolated from spleens of Tlr2^-/-^ mice (Fig. 4D). (E)-protein as well as Pam3CSK4 also enhanced secretion of IFNγ in combination with cytokines but not with TCR stimulation indicating that TCR activation is not necessary (Fig. 4E). Stimulatory effects were absent in CD8+ T cells with a deletion of TLR2 (Tlr2^-/-^) (Fig. 4E). Pam3CSK4 effects were also abolished in Tlr2^-/-^ CD8+ T cells (Fig. 4E). Using isolated blood leukocytes, we confirmed co-activation of CD69 on blood CD8+ T cells by (E)-protein (and Pam3CSK4) upon cytokine treatment which was abrogated in TLR2-deficient cells (Fig. 4F). We did not observe, however, significant co-stimulation by (E)-protein in TCR agonist-treated blood leukocytes (Fig. 4F) as these cells already showed maximal CD69 surface expression in the absence of co-stimulatory (E)-protein or Pam3CSK4 (data not shown). Our *ex vivo* data thus clearly demonstrate TLR2-dependent bystander activation of CD8+ T cells by (E) protein.

**Fig. 4.**
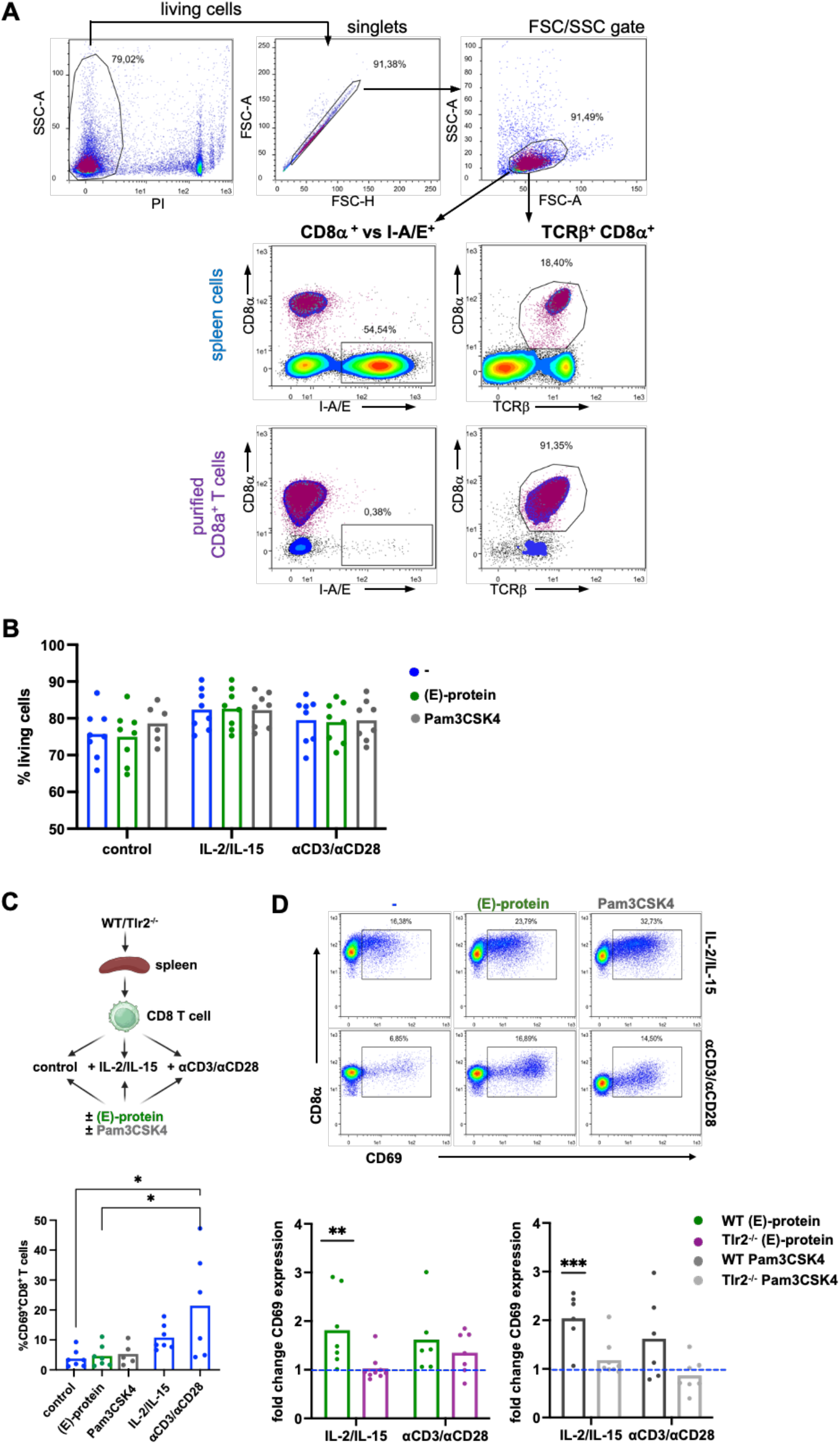
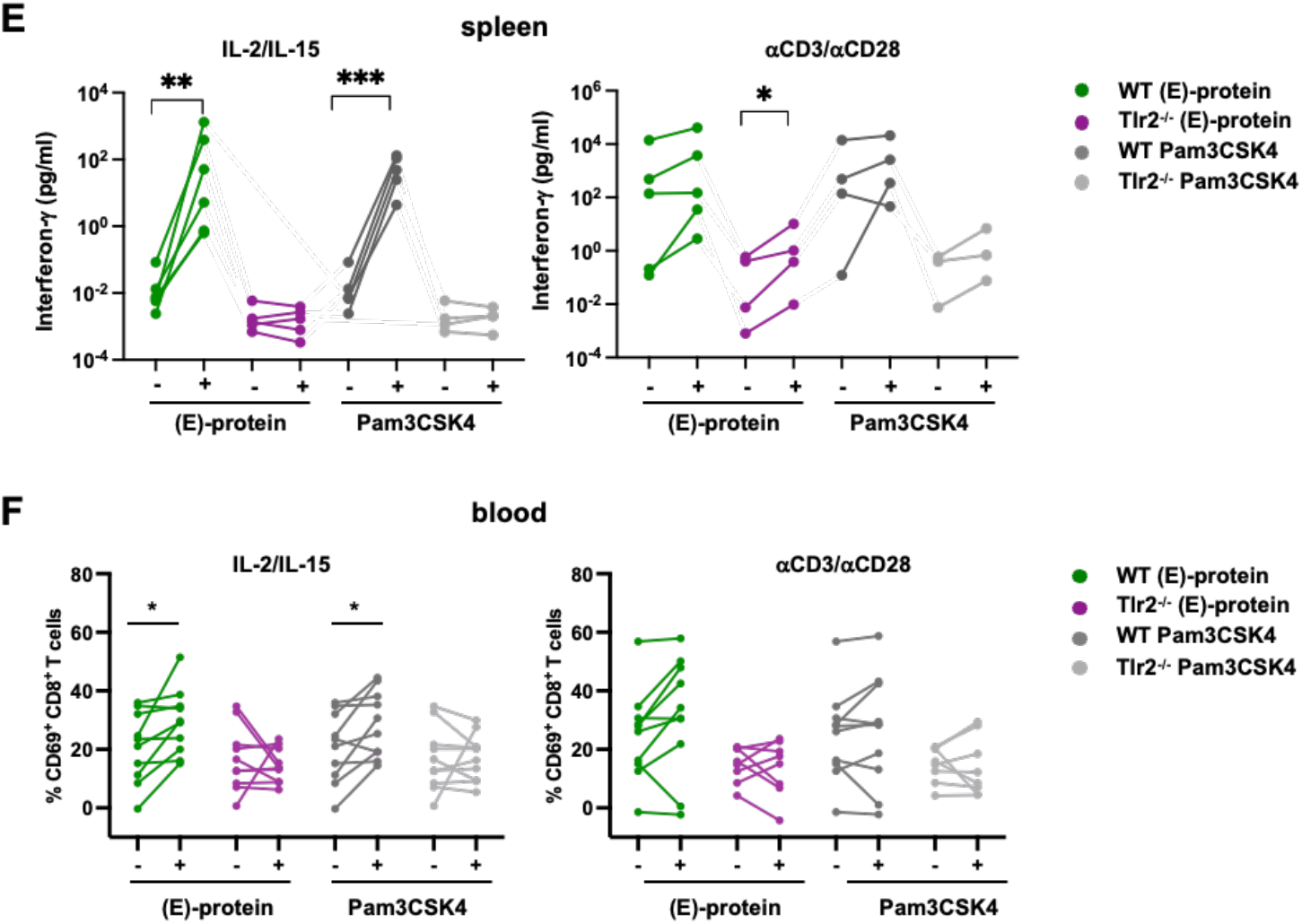
T cells are activated by (E)-protein by cytokine and antibody co-stimulation. **A)** CD8+ T cells were isolated from spleens of WT and Tlr2-/- mice using Mouse CD8α + T Cell Isolation Kit (Miltenyi Biotec) and the autoMACS Pro separator. A representative example is shown. Cells were gated for dead cell and doublet exclusion and by FSC/SSC (upper panel). Staining of spleen cells by CD8α and TCRβ specific antibodies indicates CD8+ cell population before (middle panel) and after depletion (lower panel). Staining by I-A/E specific antibodies (mouse MHC-II) indicates potential antigen presenting cells – in spleen mainly B cells - as dominant population in the sample before purification and a minor population in purified CD8+ T cells. **B)** Purified WT CD8+ T cells were cultivated without any stimulation or in presence of (E)- protein (2 μg/ml,), Pam3CSK4 (10 ng/ml), interleukin-2 (IL-2, 20 ng/ml) and IL-15 (5 ng/ml) or plate-coated antibodies αCD3 and αCD28 for 16 hr. CD8+ T cell viability was determined by flow cytometry as the percentage of propidium iodide negative cells from analyzed cells. **C)** WT spleen-isolated CD8+ T cells were cultivated either without any stimulation (control) or stimulated as above. T cell activation was determined by detecting CD69 expression on CD8+ T cells using MACSQuant Analyzer 16 and is shown as percentage of CD69+ CD8+ T cells. Pair-wise significance was calculated using Two-way ANOVA with Tukey’s multiple comparison testing, p<0.05 (*). **D)** Co-stimulatory effects of (E)-protein and Pam3CSK4 on purified CD8+ T cells. T cell activation was quantified by CD69 expression on CD8+ T cells upon cytokine (IL-2/IL-15) or TCR stimulation (αCD3/αCD28). A representative flow cytometry analysis is shown with quantification of CD69 expression as fold activation over the respective control after background subtraction (WT: n=8; Tlr2^−^/^−^: n=9). Pair-wise significance was calculated using Two-way ANOVA with Tukey’s multiple comparison testing, p<0.01 (**) and p<0.001 (***). **E)** Supernatants of stimulated WT and Tlr2-/- CD8+ T were analyzed for IFNγ and is shown in log scale. Significance was calculated using the ratio paired t-test with p<0.05 (*), p<0.01 (**) and p=0.0001 (***). **F)** Blood leukocytes were stimulated for 16 hr either with cytokines or TCR agonists in the presence or absence of (E)-protein or Pam3CSK4. Activation of CD8+ T cells was determined by CD69 surface expression after background subtraction. Starting point (cells stimulated either by cytokines or antibodies only) was the same for (E)-protein or Pam3CSK4 stimulated samples. Significance was calculated with paired t test p<0.05 (*).

### (E)-protein induced CD8+ T cell recruitment depends on TLR2 in vivo

Our *ex vivo* data prompted us to investigate the role of TLR2 in the recruitment of CD8+ T cells upon (E)-protein treatment *in vivo*. For that we applied lung-IVM in Tlr2^-/-^ mice and monitored T cell and neutrophil infiltration by the injection of fluorescently labelled antibodies 4 hours after (E)-protein instillation. (E)-protein was effectively delivered into the lungs in both, WT and Tlr2^-/-^ mice as evidenced by strong infiltration of fluorescently labelled neutrophils in both mouse strains (Fig. 5A). Deficiency of TLR2, however, completely prevented early (E)-protein-induced infiltration of CD8+ T cells into the alveolar region (Fig. 5B). CD4+ T cell infiltration was not significantly elevated upon (E)-protein instillation and not affected by the absence of TLR2 (Fig. 5B). These *in vivo* data thus strongly corroborate that the rapid infiltration that is induced by (E)-protein challenge into the lungs is mediated via TLR2 sensing. We speculate that the rapid CD8+ T cell activation by (E)-protein *in vivo* is mediated by acute cytokine signalling via acute neutrophil and macrophage activation.

**Fig. 5.**
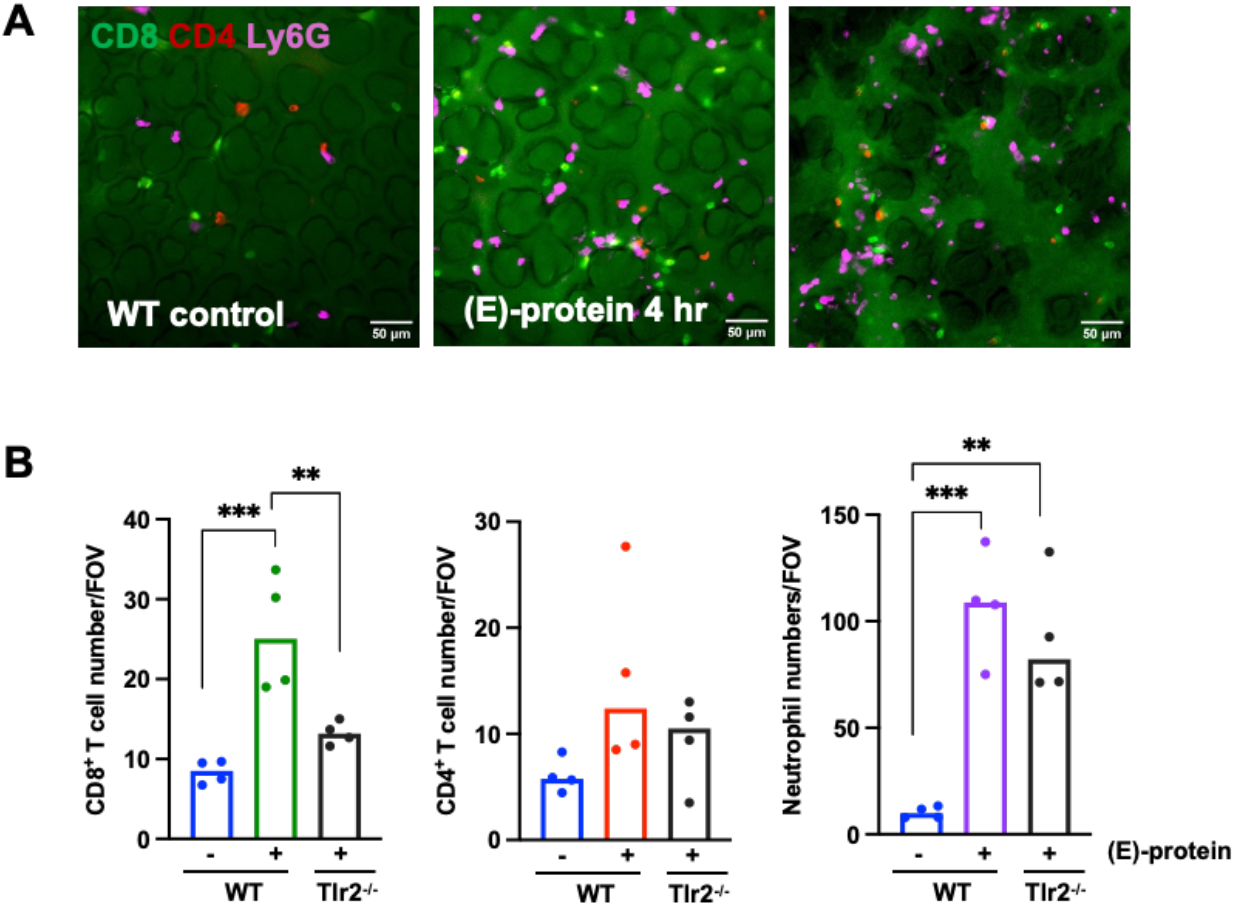
CD8+ T cell recruitment is abolished in Tlr2-/- mice. A) Representative images taken from lung-IVM time lapse videos are shown with CD8+ T cells labelled in green, CD4+ T cell labelled in red, and neutrophils detected by Ly6G labelling depicted in purple. C57Bl/6J WT or Tlr2^-/-^ mice were intratracheally instilled with or without (E)-proteins 4 hr before and intravenously injected with fluorescently labelled antibodies. **B)** CD8+ and CD4+ T cells and Ly6G+ neutrophils were quantified per Field of View (FOV, 0.16 mm^2^) using the mean of 5 FOV/mouse (n=4 mice/group). One-way ANOVA with Dunnett’s multiple comparisons test was applied with p<0.01 (**), p<0.0001 (***).

## Discussion

Antigen-specific and TCR-dependent CD8+ T cell responses are crucial for effective immune responses against SARS-CoV-2 infection. They are not only essential for viral clearance but also provide memory and cross-protection from viral variants (Moss, 2022). The differentiation of antigen-specific adaptive T effector cells is commonly assumed to require several days following antigen- and TCR-mediated priming. We here discover an early TLR2-mediated recruitment and activation of CD8+ T cells into cytotoxic T cells by the SARS-CoV2 envelope protein within 4-24 hours after (E)-protein challenge *in vivo*. Our data suggest bystander activation of CD8+ T cells and support the conclusion that (E)-protein can modulate CD8+ T cell responses during viral infection, highlighting potentially clinically relevant aspects of anti-viral response.

To investigate the mechanism underlying this rapid response, we isolated CD8+ T cells from splenocytes of naïve wild-type and Tlr2^−^/^−^ mice and assessed the stimulatory effects of (E)-protein alone and in combination with cytokines (IL-2/IL-15) or TCR agonists (αCD3/αCD28 antibodies). Neither (E)-protein nor the TLR2 agonist Pam3CSK4 were sufficient to activate CD8+ T cells on their own, indicating that residual antigen-presenting cells (APCs) present at low frequency in the purified CD8+ T cell preparations were unable to provide TCR-stimulating antigen presentation. In contrast, when combined with cytokines or TCR agonists, (E)-protein and Pam3CSK4 significantly upregulated CD69 expression and, in the case of cytokine co-stimulation, promoted IFNγ secretion, suggesting that TCR engagement is dispensable for this co-activatory effect. Moreover, stimulatory effects were abolished in Tlr2^−^/^−^ CD8+ T cells, establishing TLR2 as a necessary mediator of (E)-protein-induced CD8+ T cell co-activation. This pattern is consistent with TLR2-mediated bystander activation, a process by which innate receptor signaling amplifies CD8+ T cell responses independently of antigen-specific TCR engagement. TLR2 activation has previously been shown to control antiviral CD8+ T cell immunity in the absence of cognate antigen (Cottalorda et al., 2009; Lee et al., 2022; Whiteside et al., 2018; Zhang et al., 2019)). SARS-CoV-2 (E)-protein has been independently identified as a direct TLR2 ligand capable of activating NF-κB signaling in human monocytes and macrophages (Planès et al., 2022; Zheng et al., 2021). Hence, our *ex vivo* data suggest that (E)-protein exploits this same TLR2-dependent pathway to co-activate CD8+ T cells, cooperating with cytokine or TCR signalling in a manner yet to be fully elucidated. While we are aware that *in vitro* systems cannot fully recapitulate the *in vivo* conditions, we view these data as complementary to our *in vivo* data. The absence of responses in Tlr2^−^/^−^ CD8+ T cells supports a role for TLR2 in mediating the effects of (E)-protein. Importantly, this aligns with our *in vivo* observations of CD8+ T cell recruitment and enhanced activation. Limitations of our study include the high and probably unphysiological concentration of recombinant (E)- protein used in our *ex vivo* and *in vivo* studies. While this relative high concentration might limit the translational relevance, we argue that the combination of live lung imaging and *ex vivo* T cell analysis provides proof-of-concept data that identify a novel and so far, unknown, mechanism for viral proteins in lung infection. Moreover, we cannot fully exclude pore-forming activity of (E)-protein that has been proposed previously (W.-A. Wang et al., 2023; Zhou et al., 2023). The SARS-CoV2 (E)-protein is anchored in the lipid bilayer and self-oligomerizes to form pentameric ion channel in the cell membrane, also known as viroporin, with a selective permeability for cations such as K^+^, Na^+^ and Ca^2+^, acting as an independent virulence factor, inducing cell death and an inflammatory cytokine storm (Nieto-Torres et al., 2015). In our *ex vivo* experiments, however, we did not observe signs of cytotoxicity by the addition of (E)-protein to the purified CD8+ T cells suggesting the absence of viroporin activity in CD8+ T cells. This also applies to our *in vivo* setting where we did not observe pronounced signs of cell death aligning with previous publications on the *ex vivo* and *in vivo* application of (E)-protein (Zheng et al., 2021). We thus assume that the pore forming activity of (E)-proteins requires additional viral components for forming ion channels in ERGIC membranes that promote egress of the newly synthesized viral particles from the cell (W.-A. Wang et al., 2023). Early innate bystander CD8+ T cell activation has also been observed in SARS-CoV2 infected humans correlating with disease resolution and prevention of clinical progression to severe disease (Bergamaschi et al., 2021). We therefore concur with Karl et al. who suggested that *“…bystander-activated T cells may play an important role in the early viral defense”* (Karl et al., 2025). While TLR2 activation has been linked to COVID-19 severity (Mantovani et al., 2023) with TLR2 antagonists demonstrating protective and adjuvant effects (Mantovani et al., 2023; Zheng et al., 2021), early activation of CD8+ T cells is conversely associated with protection against severe disease (Sette et al., 2023). These results highlight that TLR2 outcomes are varied and context dependent. Further research is needed to harness our observation of a rapid SARS-CoV-2 (E)-protein–induced CD8+ T cell response into a viable diagnostic and therapeutic applications.

## Acknowledgements

We acknowledge the excellent technical assistance of David Kutsche at Helmholtz Munich and the help of the animal facilities at Helmholtz Munich and the Research Center Borstel (RCB) as well as the support by the flow cytometry unit at RCB.

